# Inferring translational heterogeneity from ribosome profiling data

**DOI:** 10.1101/866582

**Authors:** Pedro do Couto Bordignon, Sebastian Pechmann

**Affiliations:** Département de Biochimie, Université de Montréal, 2900 Blvd Edouard-Montpetit, Montréal, QC, H3T 1J4, Canada

## Abstract

Translation of messenger RNAs into proteins by the ribosome is the most important step of protein biosynthesis. Accordingly, translation is tightly controlled and heavily regulated to maintain cellular homeostasis. Ribosome profiling (Ribo-seq) has revolutionized the study of translation by revealing many of its underlying mechanisms. However, equally many aspects of translation remain mysterious, in part also due to persisting challenges in the interpretation of data obtained from Ribo-seq experiments. Here, we show that some of the variability observed in Ribo-seq data has biological origins and reflects programmed heterogeneity of translation. To systematically identify sequences that are differentially translated (DT) across mRNAs beyond what can be attributed to experimental variability, we performed a comparative analysis of Ribo-seq data from *Saccharomyces cerevisiae* and derived a consensus ribosome density profile that reflects consistent signals in individual experiments. Remarkably, the thus identified DT sequences link to mechanisms known to regulate translation elongation and are enriched in genes important for protein and organelle biosynthesis. Our results thus highlight examples of translational heterogeneity that are encoded in the genomic sequences and tuned to optimizing cellular homeostasis. More generally, our work highlights the power of Ribo-seq to understand the complexities of translation regulation.

## INTRODUCTION

Translation of messenger RNA (mRNA) into protein sequences by the ribosome is the central step of protein biosynthesis and the energetically most expensive process in the cell (*1*). Accordingly, translation is tightly controlled and heavily regulated to sustain cellular homeostasis (*2*). In turn, the dysregulation of translation is associated with many severe human diseases including cancer and neurodegeneration (*3, 4*). The kinetics of translation initiation exert the strongest influence on final protein synthesis rates (*5*). However, translation elongation, the speed and dynamics at which ribosomes move along mRNAs, is equally heavily regulated and emerging as an important aspect to defining the fate of the encoded nascent polypeptides (*6–8*).

Ribosome profiling (Ribo-seq), the experimental high-resolution measurement of translation kinetics through deep-sequencing of ribosome-protected footprints, has revolutionized the study of translation (*9–11*). Next to the accurate identification of translated sequences in genomes (*12*), many fundamental insights into the determinants of translation elongation kinetics could so far be derived from Ribo-seq data. For instance, codon translation rates on average correlate with cellular tRNA abundances (*13–15*). Positively charged amino acids can slow down translation through interaction with the negatively charged interior of the ribosome exit tunnel (*16, 17*), as can upstream RNA secondary structures that hinder ribosome translocation (*18*). Specific sequence motifs can stall translating ribosomes such as select codon pairings (*19*), or successive proline residues that require the specialized translation factor EF-P for their translation (*20*). Moreover, these determinants of translation elongation kinetics (*21–23*) are selectively placed along mRNA and protein sequences to prevent ribosome traffic jams and overall optimize translation (*24–26*).

Equally importantly, the kinetics of translation elongation directly coordinate downstream processes on the ribosome important for determining the fate of newly synthesized proteins (*6*). This includes cotranslational protein folding (*27–29*), interaction of nascent chains with chaperones or ribosome-associated targeting factors (*30–32*), protein complex assembly (*33*), and even cotranslational protein degradation (*34*). Accordingly, the tuning of translation elongation is coupled to powerful feedback systems of cellular regulation (*35*), for instance in response to stress (*36–38*).

Yet, despite tremendous progress made possible through the emergence of Ribo-seq (*9*), many aspects of translation remain mysterious. In part this must be due to the complexity of the underlying biology that awaits to be further uncovered at rapid pace. However, in part this may also stem from persisting challenges in the interpretation of Ribo-seq data that are often characterized by high intrinsic variability. For instance, the inference of individual codon translation rates from Ribo-seq data commonly results in wide distributions rather then narrowly defined individual values (*39–42*). This renders the mechanistic modelling of translation, even though strong progress could be made (*25, 41, 43–45*), challenging. Some uncertainty in Riboseq data could be directly attributed to technical difficulties and experimental biases (*14,46–48*). However, Ribo-seq experiments have been continuously improved (*49–51*) and technical noise alone cannot explain all the variability observed in Ribo-seq data.

To this end, the process of translation itself is known to exhibit substantial heterogeneity (*52*). Eukaryotic cells contain up to 10^8^ ribosomes (*53*), and variation in composition, modification, interaction, and localization can specialize their functionality (*54–56*). Such specialized ribosomes have been found for instance to prioritize the efficient translation of proteins destined for export to mitochondria (*57*). Similarly, ER-targeted proteins are preferentially translated by a select pool of translocation competent ribosomes (*58*). Heterogeneity equally exists at the level of translation elongation. The cellular abundances of tRNAs affect the effiency of diffusion to the ribosome, thus codon-specific translation rates (*59*). Long considered stable in their expression, the principles and functional implications of regulating tRNA abundances are only starting to emerge (*37*). Another mechanism that directly affects the decoding efficiency is achieved through RNA modifications at or near the tRNA stem loop (*60, 61*). Of note, tRNA modifications are well characterized biochemically but their physiological roles in most cases remain poorly understood (*60,61*). An example of a universally conserved tRNA modification is the modification of the anticodon wobble uridine (*U*_34_) in the tRNA genes *tE^UUC^*, *tK^UUU^*, and *tQ^UUG^* that increases the efficiency of translating AAA, CAA, and GAA codons (*40,62*). Taken together, a plethora of processes within the cell exists that can render translation heterogeneous, which should be reflected in Ribo-seq data.

Here, we have performed a comparative analysis of Ribo-seq data from *S. cerevisiae* to systematically detect principles of translational heterogeneity from data obtained by bulk Ribo-seq experiments. By systematically identifying subsequences whose differential translation cannot be explained solely by variability between experiments, we present cases of biological heterogeneity of translation that are encoded in the genomic sequences and likely tuned to optimizing protein and organellar biogenesis. Our results suggest that some variability commonly observed in Ribo-seq data actually represents fascinating functional biology.

## RESULTS

Ribo-seq experiments afford to probe the dynamics of translation at very high resolution through the selective sequencing of ribosome protected footprints. Nonetheless, due to the complexities of the experiment and underlying biology, Ribo-seq data published so far often contain a high degree of intrinsic variability (*48*). This was equally apparent from our comparative analysis of data from 20 independent Ribo-seq experiments comprising both biological replicas from the same laboratories as well as data obtained by different research groups (*see Methods*; Table 1). Illustrated for the exemplary yeast gene *LTV1*, we found at first sight substantial variability between the corresponding ribosome density (RD) profiles of individual experiments (Figure 1A). At the same time, many characteristic peaks of high RD that indicate regions of slow translation were strongly preserved across experiments (Figure 1A). Thus, while there was variability in these data, strong translational attenuation could clearly be detected by Ribo-seq systematically.

**Table 1:** Ribosome profiling datasets analyzed in this study.

| Publication | SRA accession | Reference |
| --- | --- | --- |
| Gardin (2014) | SRR1002819 | (98) |
| Gerashchenko (2014) | SRR1520311 | (91) |
| Guydosh (2014) | SRR1042855 | (99) |
| Pop (2014) | SRR1688545 | (100) |
| Nedialkova (2015) | SRR1944981-3 | (40) |
| Young (2015) | SRR2046309,10 | (101) |
| Lecanda (2016) | SRR3945926,8 | (50) |
| Radhakrishnan (2016) | SRR3493886,7 | (102) |
| Beaupere (2017) | SRR4000288,9 | (103) |
| Gerashchenko (2017) | SRR363557,8 | (104) |
| Schuller (2017) | SRR5008134,5 | (105) |
| Zou (2017) | SRR5090936 | (106) |

**Figure 1.**
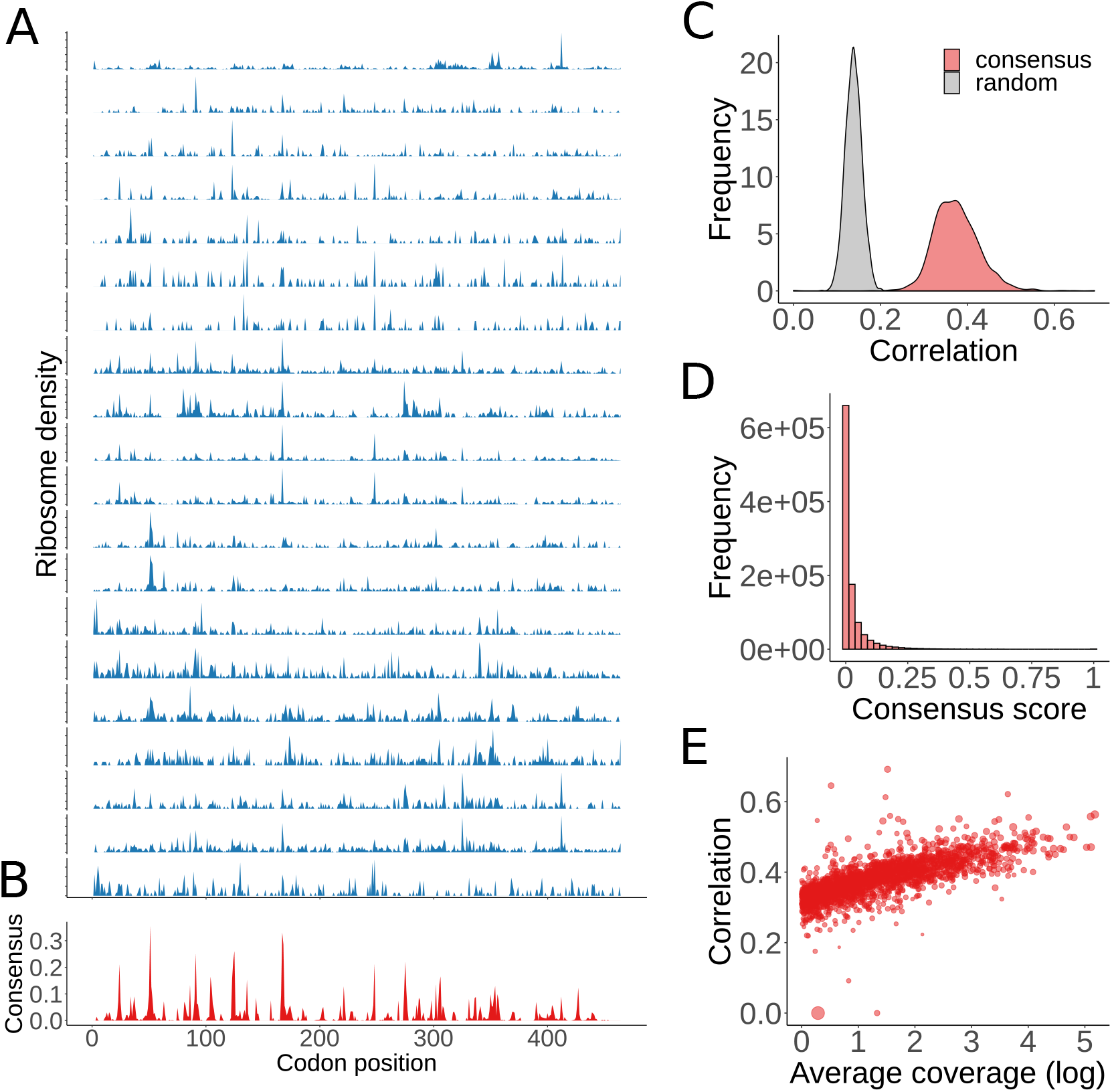
Consensus ribosome density profiles. **(A)** Ribosome density (RD) profiles from independent experiments for the exemplary yeast gene *LTV1*. **(B)** Consensus ribosome density profile for the gene *LTV1* derived from the individual experiments shown in (a). **(C)** The consensus profile is representative of individual experiments. Show are the distributions for the average correlation coefficient between individual experiments and the corresponding consensus as well as a randomized control (*see Methods*). **(D)** High consensus values are rare. Distribution of RD consensus scores across all genes analyzed. **(E)** Sequencing depth limits the resolution of ribosome profiling data. Higher sequencing coverage on average (reads per position) correlate more strongly between profiles from individual experiments and their consensus.

### Consensus RD profiles represent individual experiments

To identify a representative RD signal that is supported by independent individual experiments, we sought to derive a consensus RD profile along transcripts. Specifically, regions of high RD along mRNAs that are present in all or most individual profiles should be clearly reflected in a consensus profile as they likely represent strong signatures of translation attenuation. Due to the sensitivity of arresting translation during Ribo-seq experiments without use of cycloheximide, RD peaks in individual experiments may not be found at exactly the same positions but very slightly shifted. To account for this, we chose to derive a consensus profile upon applying a peak-synchronization algorithm (*63*) (*see Methods*). Importantly, the resulting consensus RD profile visibly reflected the main features of the individual profiles for the gene *LTV1* (Figure 1B).

To test whether our consensus generally represented the individual experiments, we computed the per-gene average of the correlation coefficients between the individual profiles and the consensus. As a control, we calculated a consensus and average correlation for sets of individual profiles randomly chosen from different genes (*see Methods*). We found much higher correlation coefficients for the true data compared to the random control, suggesting that our consensus profiles indeed well reflected the individual experiments (Figure 1C).

The obtained consensus scores ranged by definition from [0, 1], but high consensus scores were found to be very rare. Across all genes the vast majority of positions exhibited consensus scores of or just above 0, while only few positions had relatively high consensus scores (Figure 1D). As a result, peaks in the consensus profile were generally sparse and very clearly separated from the background signal (Figure 1B).

Moreover, we found that how well the consensus represented the individual profiles depended, as expected, strongly on the sequencing coverage. Genes with higher read-depth of ribosome protected footprints were clearly found with higher correlations between individual experiments and the consensus (Figure 1E). Our analyses thus suggested that a current limiting factor of Ribo-seq may be foremost insufficient sequencing depth. Taken together, the derived RD consensus profiles clearly represented individual experiments and highlighted regions whose slow translation was detected systematically by independent experiments with a strong consensus signal that clearly separated from the background.

### Consensus profiles are biologically meaningful

We next sought to evaluate whether the derived consensus profiles were also carrying biological meaning. To test this, we compared the different RD profiles to known determinants of translation elongation. Characteristic values for codon-specific ribosome densities are commonly interpreted as dwell times and used to infer codon translation rates (*64*). Accordingly, the median of the distributions of the RD values for the 61 sense codons was computed for the individual experiments, the consensus, and as a control for a simple average (mean) profile of the individual experiments (*see Methods*). More slowly translated, or nonoptimal codons usually correspond to those that have a low tRNA adaptation index (tAI) (*59*). To test whether the consensus profile was as good or better than individual profiles at identifying nonoptimal codons, we defined the bottom 25% of the sense codons with the lowest tAI as nonoptimal and evaluated the discriminative power of the inferred codon median RDs to correctly classify them.

As previously reported, individual codon RDs distributed broadly, but nonoptimal codons were found on average with higher codon consensus RDs (Figure 2A). Reciever Operating Characteristic (ROC) curves for the individual experiments, the mean profile and the consensus quantified that the consensus was indeed as good or even slightly better at classifying the nonoptimal codons (Figure 2B). Foremost, this observation validated that the consensus profiles did not introduce artificial peaks without biological meaning, but it was nice to see that they performed very well at identifying nonoptimal codons. These results were found qualitatively independent of the definition of the nonoptimal codons. Some individual experiments performed as well or even very slightly better than the consensus, suggesting variability also between the experimental data sets. Notably, the consensus method was found very powerful to condense heterogeneous data to a strong and meaningful signal.

**Figure 2.**
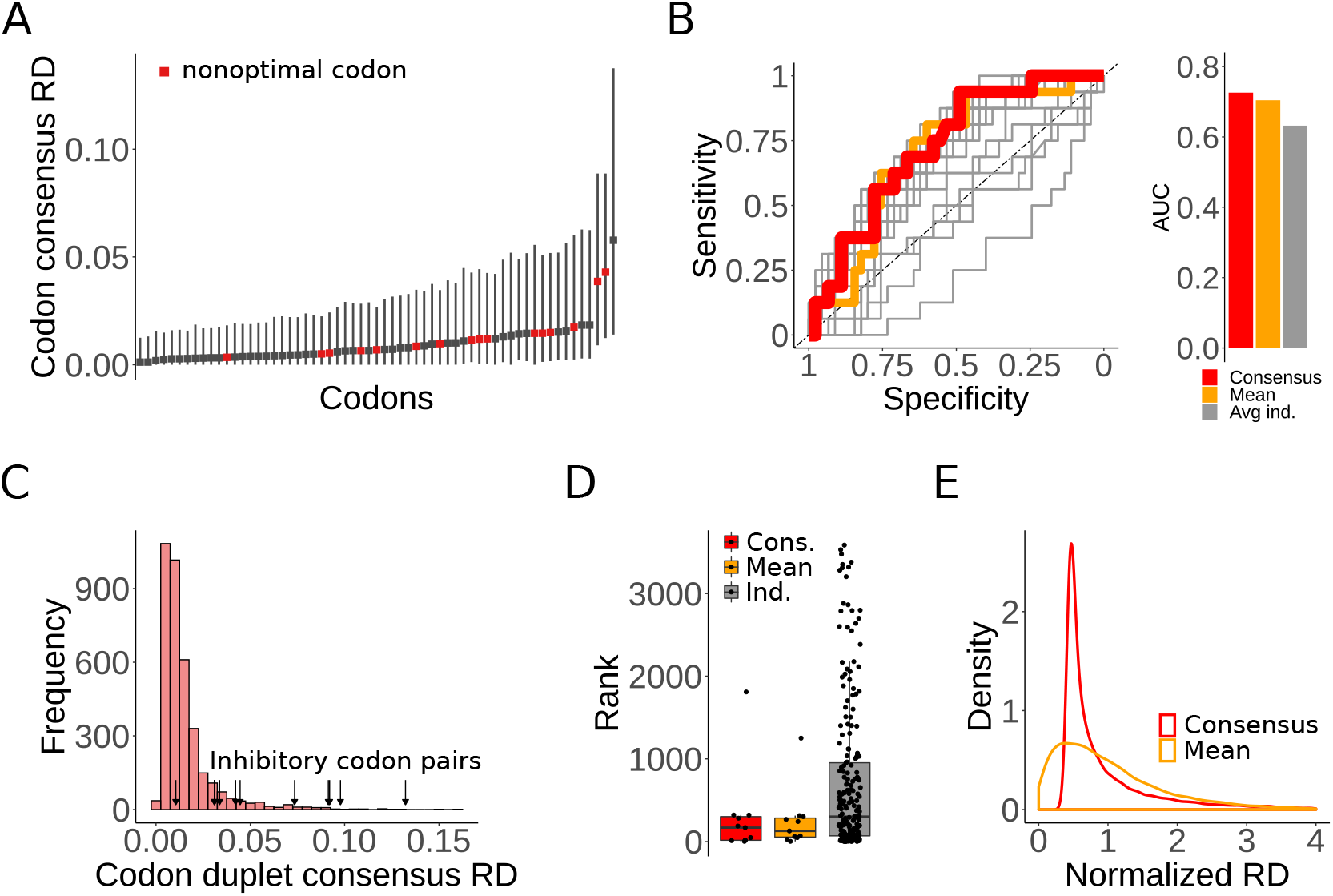
Consensus profiles identify nonoptimal codons and stalling sequences. **(A)** Distributions of consensus codon RDs for the 61 sense codons are represented by their median values (squares) and the range from 25% – 75% (solid lines). Nonoptimal codons (in red) correspond on average to higher consensus codon RDs. **(B)** The consensus profiles can identify nonoptimal codons. ROC curves for the classification of nonoptimal codons from codon RDs are shown for the consensus, mean, and individual profiles. The consensus has slightly more discriminative power than the mean or average of the individual profiles. **(C)** Inhibitory codon pairs have high consensus RD scores. Distribution of the consensus RDs of all codon pairs. Codon pairs that have been reported as strong stalling sequences are highlighted and are predominantly found in the tail of the distribution. **(D)** Rank order of inhibitory codon pairs.

Similarly, experimental work has established that select codon duplets act as very strong stalling sequences (*19*). Here, the codon duplet consensus RDs of these ‘inhibitory’ codon pairs (*19*) were almost all found to be much higher than average (Figure 2C). Of note, we found inhibitory codon pairs strongly selected against in the set of analyzed genes, reflected by only very few occurrences in yeast mRNA sequences compared to the numbers of other codon pairs. Codon duplet RDs derived from both the consensus and the mean profiles ranked generally higher than in the individual experiments when sorting all codon duplet scores in ascending order (Figure 2D). These observations further underlined the power of a joint analysis of multiple ribosome profiling datasets through a consensus profile to filter out a consistent signal from heterogenous data sources.

Last, while both the consensus and the simple average of the individual profiles gave an improvement with respect to the prediction and classification of nonoptimal codons and inhibitory codon pairs, they clearly differed in how they distributed. The distribution of mean RD values followed a initially concave upper tail while the distribution of consensus RDs showed a strongly convex upper tail (Figure 2E). As a result, the high RD peaks in the consensus profile general better separated from the background signal of low RD values.

### Differentially translated (DT) sequences are rare

Individual codon RDs obtained from the consensus profiles displayed broad distributions similar to what had been reported previously (*39–42*). This observation suggested substantial variation of the speeds at which individual codons are translated. Some of these differences may simply be due to intrinsic noise in the complex sequencing experiment. However, if the same codons, or subsequences of codons, could be found differentially translated beyond what can be accredited to experimental variability, then the observed differences may also have biological origins that reflect cases of translational heterogeneity.

To identify such differentially translated (DT) sequences, we first devised a randomization procedure that accounted for the gene-specific biases in coverage. Specifically, the number of peaks in the consensus profile strongly depended on the overall coverage and number of stalling sites in the individual profiles. Therefore, we computed the consensus of shuffled individual profiles for each gene to estimate expected background distributions of consensus scores (*see Methods*). At the lower end, the representative 5, 10, and 20 percentiles of expected consensus scores across genes fell into narrow distributions around very low values. This matched the observation that most positions along transcripts did not have a pronounced consensus signal and that stalling sites in general were sparse. In contrast, the representative 80, 90, and 95 percentiles of the expected consensus scores formed broad distributions, highlighting a strong dependency on gene-specific profiles (Figure 3A).

**Figure 3.**
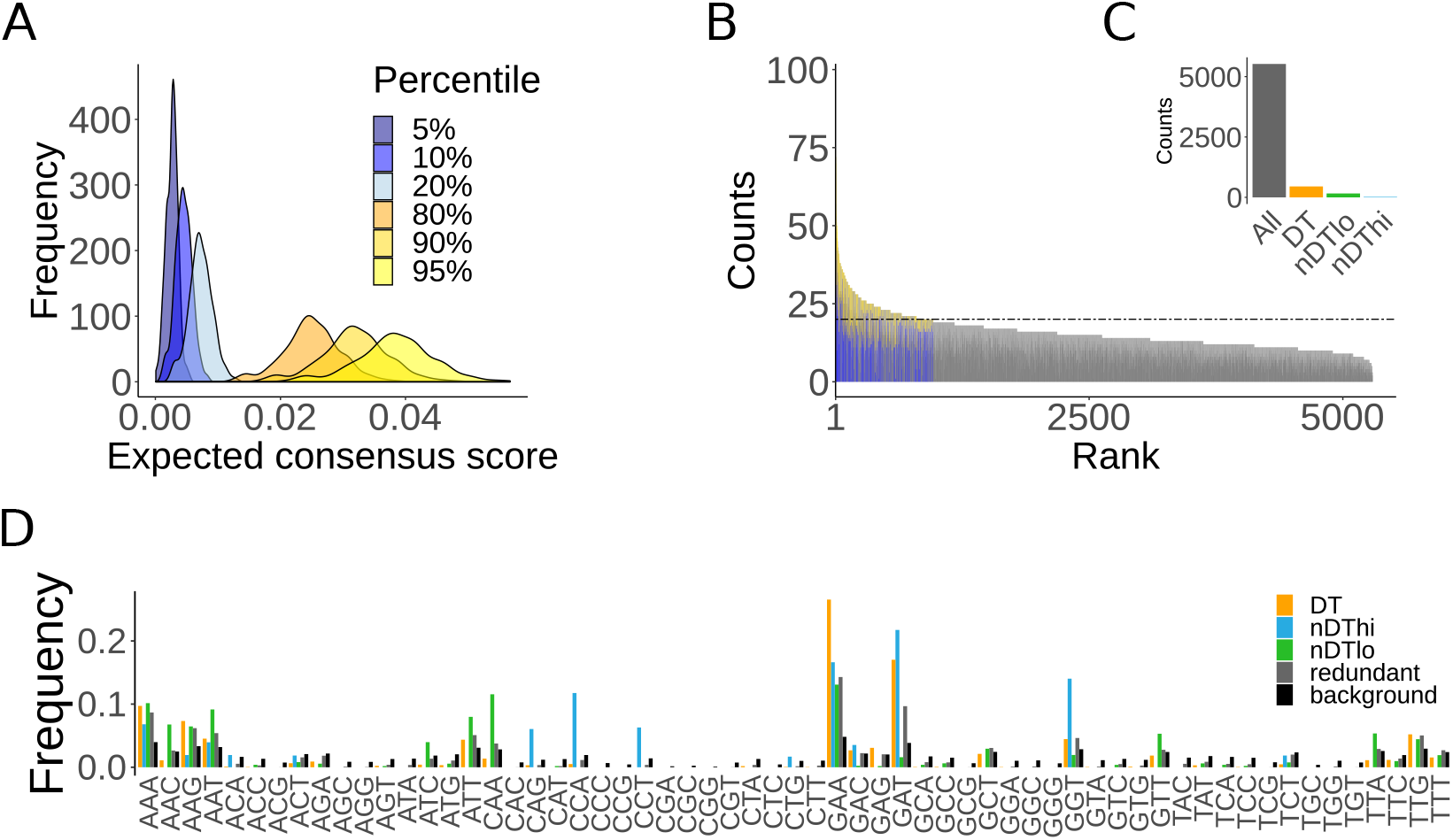
Identification of differentially translated (DT) subsequences. **(a)** Characteristic percentiles of the distributions of expected consensus scores obtained from randomization. Distributions of expected consensus scores are strongly gene-specific as apparent from broad distributions, especially at high percentiles. **(b)** Redundant subsequences and DT sequences. **(c)** Enrichment and depletion of specific codons from DT sequence.

Next, we set out to systematically identify DT sequences in the *S.cerevisiae* genome. Focussing on subsequences of length 3 codons, we defined DT sequences as those that could be found at least 10 times at or below the lower gene-specific 10% threshold and at least 10 times above the gene-specific 90% threshold (Figure 3B). Of note, the analyzed 2443 *S.cerevisiae* mRNA sequences contained 5525 subsequences of 3 codons with a general redundancy of more than 20 occurrences. The median number of occurrences of these 5525 sequences was 27, and only the top 10% were found to have a redundancy of over 44 counts. Our definition of DT sequences was therefore sufficiently stringent and selective. Importantly, of these 5525 codon triplets, 451 fulfilled our definition as DT sequences (Figure 3C). While overall the number of subsequences outside the chosen significance thresholds correlated with their overall redundancy, the observed number of DT sequences was sparse, leaving the possibility that they serve specific functions. For comparison, only 159 sequences were found as translated consistently fast (nDTlo), and only 17 as translated consistently slowly (nDThi) (*see Methods*).

We next asked whether the DT sequences were enriched in specific codons. Strikingly, the codons the most over-represented in the DT sequences were those recognized by tRNAs whose known modifications at the wobble-uridine position modulate the speed of translation, primarily sequences containing GAA codons (Figure 3C). Equally importantly, absent were codons encoding Prolines that are known to elicit strong stalling and likely leave little room for contextual modulation of translation speeds. In contrast, nDTlo sequence were enriched in CAA codons, another codon linked to the *U*_34_ tRNA modification, while nDThi sequence showed a high frequency of the Proline codons CCA and CCT that act as known strong stalling signals. Equally remarkably, when normalizing amino acid usage by transcript abundance, i.e. how often the corresponding codons are translated by the ribosome, Glutamine residues become one of the most heavily used amino acid (*27*). As such, choice between CAA and CAG codons encoding Gln residues may strongly impact overall translation efficiency. Accordingly, we observed a clear preference for the optimal codon CAA, whose decoding efficiency is equally further increased by tRNA modification, in the nDTlo sequence, and of the nonoptimal codon CAG in the nDThi sequences. In addition to distinct codon preferences, low RD DT sequences were on average found later along mRNA sequences and thus may also reflect know biases in decreasing coverage along transcripts. Taken together, our analysis suggested that DT sequences exhibited distinct codon usage patterns that strongly linked to known determinants of modulating translation elongation kinetics.

### DT sequences may be important for protein biogenesis

Having identified select sequences that are translated differentially in systematic fashion beyond what can readily be attributed to variability between experiments, we next sought to better understand what characterizes whether the same subsequence is translated slowly or fast. Clusters of several adjacent positions of low, and especially of high RD can have a more pronounced impact on translation elongation compared to individual positions. Regions of low RD can be under selection for fast translation, but can also result from limited sequencing coverage in the Ribo-seq experiments. In contrast, high RD regions generally exert a stronger influence on translation elongation and coupled processes. We therefore focused on regions of high RD and first asked whether high RD DT sequence preferentially occur in clusters of other high RD positions or not. Across the set of analyzed mRNA sequence positions of high consensus RD generally did not show a preference for being in clusters, here defined as at least 3 sites with RDs above the gene-specific upper threshold in a window of 7 positions (Figure 4A). In contrast, the vast majority of DT sequences were found to occur in clusters (Figure 4A). This observation suggested that DT sequences may contribute to the selective and potentially functional attenuation of translation.

**Figure 4.**
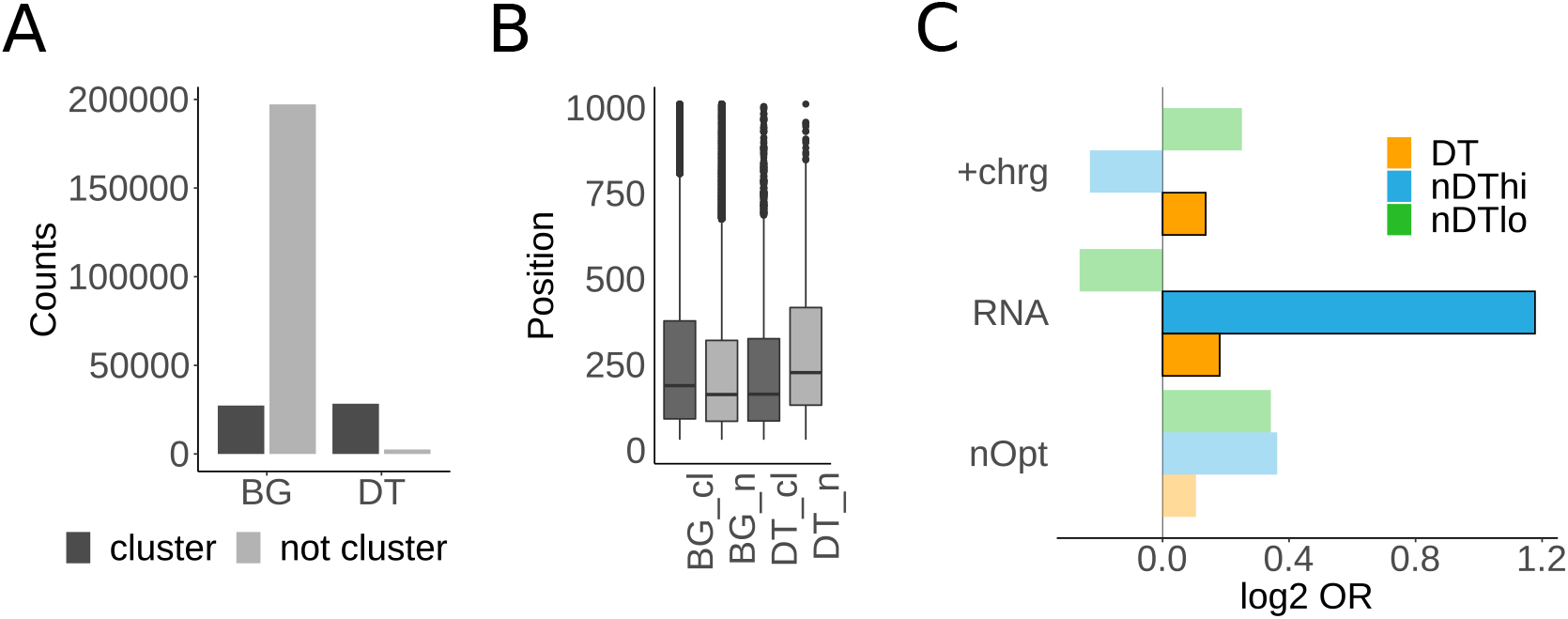
Characteristics of DT sequences. **(a)** DT sequences are preferentially found in clusters. While only a small fraction of non-redundant high consensus RD regions (BG for background) cluster, the vast majority of DT sequences with high consensus RD appear in clusters. **(b)** DT sequences in clusters (DT c) are found on average closer to the N-terminus in proteins compared to BG, and in contrast to BG earlier in sequence compared to DT sequeces not in clusters (DT n). **(c)** Known stalling signals contribute to DT sequences. Statistical associations between high consensus RD occurences of DT sequences and RNA folding strength, nonoptimal codons, and clusters of positively charged amino acids.

Moreover, we found a bias of DT sequences in clusters to be positioned earlier in sequence than those not in clusters, and in contrast to the background signal of all high consensus RD positions (Figure 4B). If translation elongation influenced protein synthesis, folding, and homeostasis, any cues to optimize translation or promote downstream processes would be expected earlier in sequence. Guiding the formation of a folding nucleus, coordinate chaperone binding, or, equally importantly, regulating ribosome spacing to prevent ribosome traffic jams, thus facilitating efficient overall translation, are all decisive events for the fate of a nascent polypeptide that usually occur early on during translation. Our result of a positional bias thus left open the possibility that DT sequences may play a role in regulating protein biogenesis.

To better understand how the sequence context may determine whether instances of the same sequence were sometimes translated fast and sometimes slowly, we analyzed a putative link to three known stalling signals, clusters of positively charged amino acids, high RNA folding strength, and clusters of nonoptimal codons. In comparison to the DT sequences that were defined as sequences with at least 10 occurrences each below and above the significance thresholds, we analyzed sequences that were consistently translated fast or slowly, i.e. non-differentially translated sequences of consistently high RD (nDThi) and low RD (nDTlo) respectively (*see Methods*). Positively charged amino acids can slow down translation through interaction with the interior of the ribosome exit tunnel. Our analysis suggested that high RD DT sequences were significantly (p = 0.007 by Fisher’s Exact test) enriched in downstream clusters of positive charges (Figure 4C). Similarly, high RD DT sequences were significantly enriched in high RNA folding strength upstream (p = 0.002 by Fisher’s Exact test), which may hinder ribosome translocation along the mRNA, thus slowing down translation (Figure 4C). Remarkably, nDThi squences, i.e. those that are consistently translated more slowly, linked to a very strong enrichment of high RNA folding strength (p = 0.03 by Fisher’s Exact test) (Figure 4C). In all cases we observed a higher number of clusters of nonoptimal codons just upstream of the translated position of high RD, albeit without significance. Taken together, these results suggested that known stalling signals play a clear role in the observed heterogeneity of translating the same sequences.

### No systematic role of DT sequences in protein folding

Based on the findings that the rhythm of translation elongation can coordinate protein folding events at the ribosome (*27–29*), we next asked whether DT sequences may systematically link to two major events in cotranslational protein folding, the binding of the main cytosolic Hsp70 chaperone SSB, and the folding of protein domains. It has been previously shown that a local slowdown of translation can coordinate the binding of chaperones or targeting factors to nascent polypeptides at the ribosome (*30, 32*). Once bound to an elongating nascent chain, a chaperone can fundamentally change the folding landscape of the protein. Therefore, a coordinated attenuation of translation may increase the likelihood of chaperone binding to promote correct folding. We observed, as previously reported, a distinct and localized higher average consensus RD in the translated mRNAs just when Hsp70 is binding to the elongating nascent polypeptide outside the ribosome exit tunnel (Figure 5A). Notably, this signal is much more pronounced when only considering the first binding site per protein rather than all of them. However, we could not observe any localized enrichment of DT sequences within the same distance of Hsp70 binding sites (Figure 5A). This result reinforced that local translation kinetics may contribute to coordinating the binding of cotranslationally acting chaperones, but suggested that this is independent of DT sequences.

**Figure 5.**
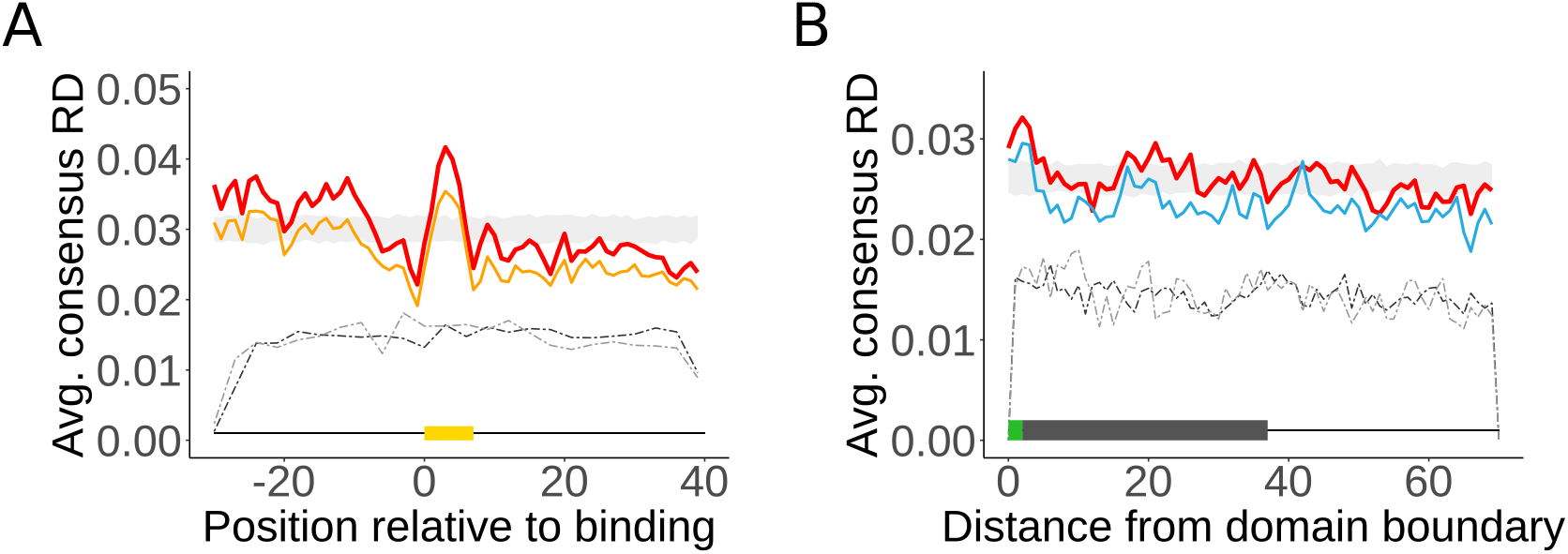
DT sequences are not systematically coordinating protein folding on the ribosome. **(a)** Meta-gene analysis Hsp70 SSB binding sites. Reported are per-position average consensus RD profiles of mRNA regions that are translated while Hsp70 is binding cotranslationally to the encoded elongating nascent polypeptide outside the ribosome exit tunnel; shown are profiles for the first Hsp70 binding site in each substrate protein (red line) and all Hsp70 binding sites (orange line) relative to a randomized background indicating *mean ± SD* (grey ribbon). On average, translation slows down when Hsp70 is binding (indicated by yellow area), as observed by (*32*). The density of high (light grey dashed line) and low (dark grey dashed line) DT sequences shows no elevated occurrence that could contribute to this local translation attenuation. **(b)** Meta-gene analysis of protein domain boundaries. Reported are per-position average profiles consensus RD downstream of the first domain boundary (green area) per multi-domain protein (red line), and for only domain boundaries that precede any Hsp70 chaperone binding (blue line). No systematic translation attenuation linked to domain folding could be observed for the consensus RD values, although a local slowdown of translation may coordinate the folding of some protein domains that precede Hsp70 binding (blue line) outside the ribosome exit tunnel (grey area). The density of high (light grey dashed line) and low (dark grey dashed line) DT sequences shows no elevated occurrence that could contribute to local translation attenuation.

Similarly, it had been previously observed that a local attenuation of translation is linked to protein domain boundaries, leaving additional time for a completely translated and exposed domain to fold outside the ribosome before translation continues at high speed (*65*). However, this could so far only be observed for individual proteins, not systematically (*66*). Similarly, we could not detect any systematic local slowdown of translation that may link to internal protein domain boundaries (Figure 5B). Accordingly, we found no enrichment of DT sequences that may coordinate protein domain folding (Figure 5B).

Thus, DT sequences appeared to be neither coordinating Hsp70 binding or domain folding in a systematic fashion. Because these are pivotal events in *de novo* protein folding, any programmed translation slowdown may not be encoded through DT sequences but rather through other, more unambiguous stalling signals. Our results leave open the possibility that DT sequences are involved in the dynamically regulated coordination of such protein folding events in individual cases that however will have to be understood at much higher resolution.

### DT sequences link to protein and organelle biogenesis

Our findings that DT sequences preferentially occurred in clusters and early in sequence indicated the potential for a functional role in coordinating early protein folding events at the ribosome. However, we could not find any systematic link between DT sequences and the binding of the main cytosolic Hsp70 SSB or the folding of protein domains. Next to the direct coordination of protein folding, which will have to be understood at higher resolution of the individual folding contexts, clusters of slowly or fast translated sequence regions can also have a strong cumulative impact on the overall translational efficiency, thus final protein levels. In addition to just more efficient overall translation, clusters of slowly translated codons early in sequence can contribute to the careful spacing of ribosomes along mRNA sequences to prevent ribosome traffic jams (*24–26*).

To test whether genes enriched in DT sequences linked to specific functions within the cell, we performed Gene Ontology (GO) analyses (Figure 6). Specifically, we compared the functional annotations of the set of ‘redundant’ genes, i.e. mRNAs with the overall highest density of redundant subsequences, to five categories of genes enriched in DT sequences. Herein, we considered ‘DT’ genes as the mRNAs with the highest density of DT sequences outside the gene-specific thresholds (*see Methods*); in comparison, ‘DTmod hi’ and ‘DTmod lo’ were defined as the sets of genes with the largest difference between high and low RD DT sequences that contain GAA codons, i.e. the most overrepresented codon that is recognized by a tRNA subject to wobble-uridine modification; similarly, we considered ‘DTnmod hi’ and ‘DTnmod lo’ as the set of genes with the largest difference between high and low RD DT sequences not containing GAA codons (Figure 3D). These four additional lists of genes thus reflected mRNAs that contained DT sequence with strong bias towards consistently fast or slowly translated, and stratified by a possible effect of the wobble-uridine tRNA modification.

**Figure 6.**
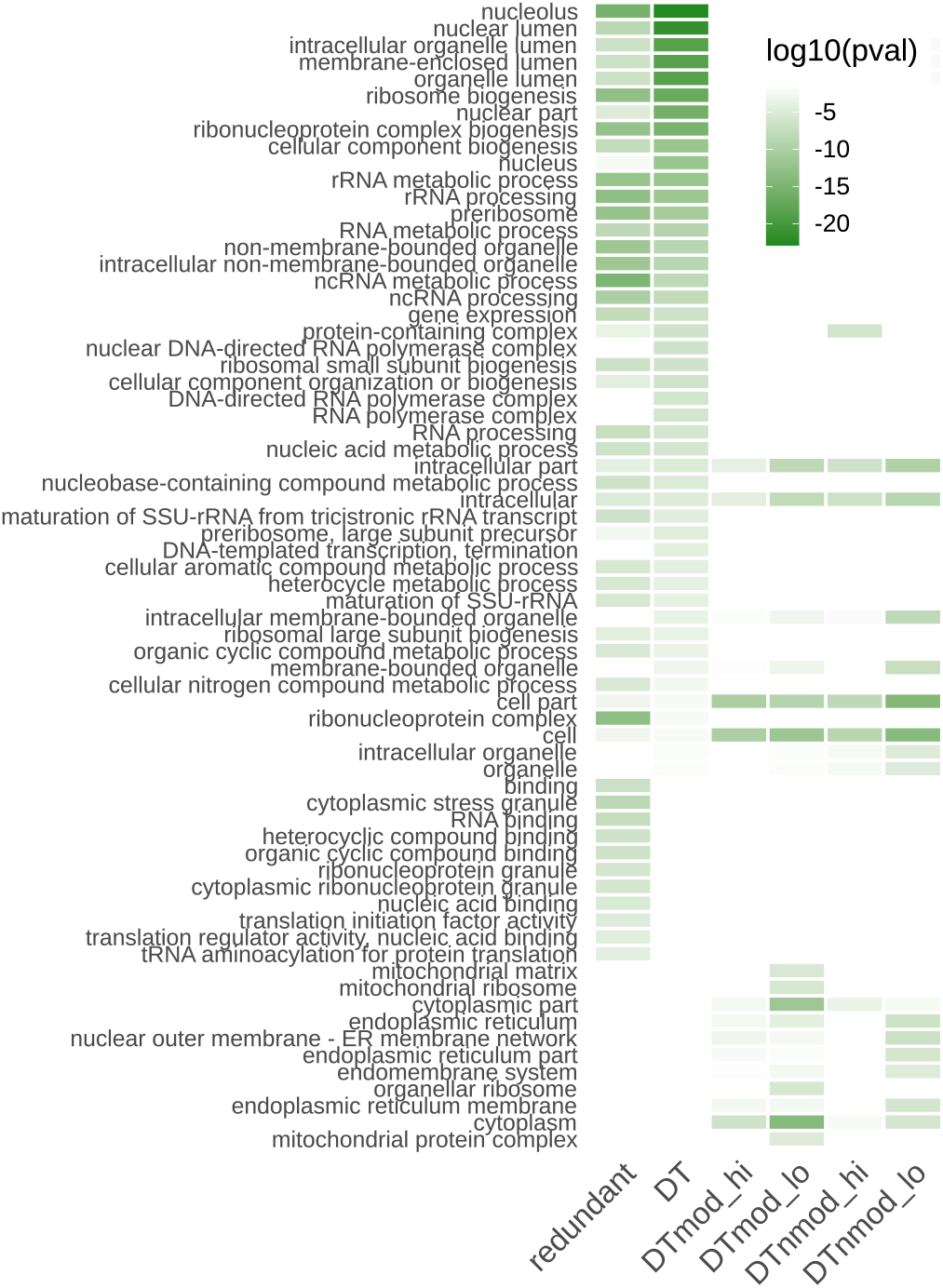
DT sequences link to protein and organelle biosynthesis. Gene Ontology (GO) analysis of DT sequences. Enrichment of GO categories color-coded base on adjusted p-value as provided by the GProfiler webserver. ‘Redundant’ denotes the top 5% of genes with the highest density of sequences of 3 codons that have at least 20 occurrences across the analyzed mRNAs; ‘DT’ here are the top 5% of genes with the highest density of DT sequences. ‘DTmod high’ and ‘DTmod low’ are the top 5% genes with the largest difference between DT sequences of high and low consensus RD respectively and that contain GAA codons recognized by tRNAs subject to modification of the anticodon wobble uridine; ‘DTnmod high’, and ‘DTnmod low’ are similarly defined for sequences that do not contain GAA codons.

Several important observations could be made. First, each category corresponded to clearly defined GO categories. This suggested that these sets of genes not only shared distinct sequence characteristics, but also functional roles (Figure 6). A general correlation between the ‘redundant’ and ‘DT’ categories was expected as a higher level of redundancy on average and by our definition also leads to higher counts of DT sequences outside the significance thresholds. Both categories were strongly enriched in genes involved in nuclear biology, RNA processing, and ribosome biogenesis. The main difference between these two categories was that the set of ‘redundant’ genes was also strongly enriched in GO categories linked to stress granule formation. Redundant or repetitive sequences are known to link to phase transition and stress granule formation (*67, 68*). In contrast, the ‘DT’ genes showed a stronger enrichment in GO categories linked to nucleus, nucleolus, and ribosome biogenesis biology (Figure 6), i.e. categories directly linked to protein biosynthesis.

Remarkably, the remaining lists of genes representing mRNAs with DT sequences that were almost only fast (low RD) or slowly (high RD) translated associated with an orthogonal set of GO terms, namely mitochondrial and endoplasmic reticulum (ER) biology (Figure 6). These included severel GO terms describing processes of organelle biosynthesis, from ‘endoplasmic reticulum membrane network’ to ‘mitochondrial matrix’ to ‘mitochondrial ribosome’ (Figure 6). In general, more functional categories associated to the low than the high RD categories. The main difference between DT sequences that included codons whose translation can be modulated by wobble-uridine tRNA modification (*40, 62*) was found to be an additional enrichment in the ‘mitochondrial ribosome’ and ‘organellar ribosome’ GO categories, as well as a link to ‘mitochondial matrix’ and ‘mitochondrial protein complex’ GO terms. Strikingly, of the genes with consistently high consensus RD, DT sequences that depended on the tRNA modification were found predominantly in mRNAs important for endoplasmic reticulum biogensis, while genes with consistently low consensus RD of their DT sequences subject to tRNA modification associated with mitochondrial biology. Similar results could be obtained by considering all three codons (AAA, CAA, and GAA) whose translation is modulated by this tRNA modification (*40, 62*). Moreover, while this tRNA modification is normally pervasive and applied to most corresponding cytoplasmic tRNAs (*60*), it may be less present in cases of localized translation known to facilitate the synthesis of proteins destined to the major organelles. Of note, mitochondrial proteins often fold in the cytosol and their export to the mitochondria often share components of the cytoplasmic protein quality control network (*69–71*), in *S.cerevisiae* for instance the Hsp70 chaperone SSA1 (*72*). In contrast, translocation to the ER is directly coupled to translation as ER-destined proteins usually cannot fold inside the cytoplasm. Moreover, ER-destined proteins are often translated by a specialized pool of translocation-competent ribosomes near the ER membrane (*58*). Our results thus suggest that the translational heterogeneity observed may be directly coupled to the organization of proteins to the major organelles, and indicate that tRNA modifications could play a role in this.

## DISCUSSION

Data from Ribo-seq experiments have lead to many fundamental discoveries of mechanisms that govern the kinetics of translation. However, challenges persist in the analysis and interpretation of Ribo-seq data that are often characterized by intrinsic variability. Here, we show that some of this variability represents cases of biological heterogeneity of translation rather than experimental biases or noise. Through a comparative analysis of published *S.cerevisiae* Ribo-seq data sets we have systematically identified short subsequences that exhibit substantial variation in how they are translated. Importantly, whether the same sequence was translated slowly or fast associated with known stalling signatures of translation elongation, including high RNA folding strength and positively charged amino acids, or linked to mechanisms that can dynamically regulation translation kinetics such as tRNA modifications. Moreover, these DT sequences were found selectively enriched in genes central to protein and organelle biosynthesis. Our results thus indicate how translational heterogeneity that is directly encoded in the genomic sequences may serve to optimize overall protein biosynthesis and cellular homeostasis.

Many aspects of translational heterogeneity, especially of localized translation or translation on specialized ribosomes, have been under increasingly intense investigation. A spectacular example illustrating the importance and immediate functional consequences of local protein biosynthesis is given by neurons where efficient protein production in distal axons is required to control synaptic transmission (*73, 74*). Similarly, the importance of localized translation in the production of proteins that constitute the major organelles is generally well established (*58,69*). Interestingly, while the translation of ER-destined proteins is almost completely separated from the production of cytosolic proteins, substantial overlap exists between the protein biosynthesis and quality control pathways for mitochondrial and cytoplasmic proteins. Accordingly, cytosolic and mitochondrial translation are tightly coordinated (*75*). The use of DT sequences, i.e. sequences that are differentially translated in a context-dependent manner, open an intriguing window into additional mechanisms that may contribute to cellular organization through translation regulation.

One such central regulatory mechanism that remains under dynamic cellular control is given by tRNA modifications, which, for instance, have been shown to play an important role in the global regulation of protein levels in response to stress (*76–79*). We found that consistently slowly translated DT sequences containing codons decoded by tRNAs subject to wobble-uridine modification, i.e. suggesting an absence of the modification, are linked to the ER. In contrast, consistently fast translated sequences containing codons recognized by tRNAs amenable by wobble-uridine modification, which is normally present in cytosolic translation, associated with genes invo lved in mitochondrial biology. Remarkably, this overall link between the biased translation of DT sequences and organellar biology is directly following the established differences in the biosynthesis of mitochondrial and ER proteins. While an intriguing observation, our work cannot currently resolve whether our result is cause or consequence: does localized translation in a different cellular milieu merely differentially predispose to the availability of this tRNA modification? Or does the differential translation regulated through absence or presence of a tRNA modification aid in the targeting of the elongating nascent chains to their correct destination? While these questions warrant further investigation, our results already highlight a fundamental conundrum: biological sequences are initially invariant, but the cellular system is highly dynamic. Robust control could easily be achieved by using unique sequences decoding slow or fast translation. Given the complexity of the cell as a system, it is most likely that the combinatorics achieved through hierarchical layers of regulation are needed to ultimately sustain homeostasis. Herein, the use of redundant sequences can pose several distinct advantages. A synchronization of the dependency on cellular resources, e.g. specific amino acids that need to be attached to the elongating nascent chain, also render these sets of genes susceptible to shared regulatory pathways. Moreover, shared sequences at both the RNA and protein levels can serve as recognition and binding sites. A well characterized example that is supported by our results is given by stress granules which are known to depend on low complexity and repetitive sequences for their formation (*67, 68, 80, 81*). Similarly, the strong enrichment of DT sequences in genes involved in ribosome biogenesis may be strongly tied to an autocatalytic control of ribosome biogenesis (*82*). Taken together, our results suggest a use of DT sequences that may enable selective feedback in regulating and optimizing protein biosynthesis.

Extensively discussed in the context of gene expression (*83*), our results also highlight a pervasive stochastic character of translation. Many of the steps required for translation elongation are stochastic, foremost the diffusion of charged tRNA-ternary complexes to the ribosome that can strongly influence decoding efficiencies. Next to the randomness of biomolecular interactions, some stochasticity may actually be built into the system. The cell has to operate efficiently under many different conditions, and this is especially true for translation as the energetically most expensive process. Translation under non-stress conditions may be not completely optimized for overall protein production efficiency, but rather to maintain agility and rapidly adapt to changing and more challenging conditions. This has been directly observed for instance under conditions of amino acid starvation that rapidly affect codon-specific translation rates (*84, 85*). More generally, the suppression of stochasticity in biological systems is very expansive and subject to cost-benefit trade-offs that usually tolerate some level of stochasticity (*86*). Thus, for modeling translation as well as predicting the effect of translation elongation kinetics on cellular processes, it may become equally important to explicitly consider a stochastic character of translation than to improve on more narrowly defined estimates of codon-specific translation rates.

Finally, our results foremost underline the power of Ribo-seq to discover fundamental aspects of translation. A main limiting factor in current Ribo-seq studies may simply be too shallow sequencing depth. While it has been suggested that inference from individual experiments may be limited (*87*), our work of deriving and analyzing a consensus signal demonstrates that a comparative meta-analysis of several datasets can yield important additional insights. Many details of programmed translational heterogeneity result from the complex interplay of many different contributions and await to be characterized at much higher resolution. Next to a better understanding of the sequence determinants of translational heterogeneity, this includes better understanding the functional roles of central tRNA modifications (*88*), rRNA modifications (*89*), and the interplay between tRNA and epigenetic mRNA modifications (*90*). Our results on identifying principles of translational heterogeneity from bulk Ribo-seq data suggest a shift in the interpretation of variability in these data and open a fascinating window into further understanding the intricacies of translation, one of the most central and energetically the most costly process in the cell. What has been considered unwanted variability in Ribo-seq data in part actually reflects exciting biology.

## MATERIALS AND METHODS

### Code and data availability

Computer code and data to reproduce all presented results is available at www.github.com/pechmannlab/riboconsensus

### Data sources and processing

We analyzed the ribosome profiling datasets from the experiments performed without use of the translation inhibitor cycloheximide (*91, 92*) that are listed in Table 1. Sequencing adapters were removed with Cutadapt (*93*). Reads that subsequently did not align to yeast ribosomal RNA or transfer RNA sequences were mapped to the *S.cerevisiae* reference genome R64-1-1 (*94*) with STAR (*95*). Ribosome density (RD) profiles of unambiguously mapped reads were computed with Scikit-ribo (*96*). Genes with an average coverage of less than 1 read per position across datasets, or with more than 30% multi-mapping, i.e. the fraction of mRNA sequences that yields ambiguous read mapping (*97*), were omitted. This yielded a final dataset of 2443 yeast genes. Furthermore, the first and last 20 codons from each gene were excluded from our analyses of translation elongation as they reflect known biases that link to translation initiation and termination (*45*).

### Consensus RD profiles

Consensus ribosome density profiles were computed with a peak synchronization algorithm initially developed for the analysis of EEG data (*63*). Herein, peaks in individual profiles were detected by segmentation above a threshold of profile *mean* + 1*SD* (standard deviation). Peaks across individual profiles were then averaged upon applying a Gaussian weight relative to their distances from the current position; in this way peaks that were systematically present at the same position yielded a maximum consensus signal, while peaks that were shifted made a reduced contribution with their weight reflecting their distance. Gaussian weights were inferred based on a probability density function with threshold *th* = 0.0001 after which tails were considered insignificant and omitted, a central coefficient *ca* = 0.5 describing the bin width at which the maximum weight was applied, and *SD* = 1. The resulting consensus profiles were found very robust to the choice of these parameters as previously reported (*63*). Multi-mapping positions in sequences were excluded from computing the consensus profile. The average correlation to consensus was computed as the per-gene average of the pairwise Spearman correlation coefficients between individual experiments and the corresponding consensus profile. The randomized control was generated by computing a consensus profile of individual experiments randomly drawn from different genes with a minimum length of 100 codons. Similarly, per-gene background percentiles were derived from computing the consensus from shuffled individual profiles of the same gene 100 times and analyzing the distribution of the resulting consensus values. In both cases multi-mapping positions were omitted. For comparison, a simple average profile was computed as the per-position average of the individual profiles normalized to *mean* = 1.

### Validation of consensus profiles

To validate that the consensus RD profiles carried biological meaning, we sought to evaluate their predictive power for known stalling signals, notably nonoptimal codons and inhibitory codon pairs. The RDs of individual codons are commonly interpreted as ribosome dwell times and used to infer codon specific translation rates. Because Scikit-ribo accurately maps the translated codon in the ribosome A-site (*96*), we used the Scikit-ribo output to infer characteristic codon RDs as the median of the distribution of RDs of all occurrences for each sense codon or codon pair. The first and last 20 codons from each gene and multi-mapping positions were omitted. Individual profiles were normalized to *mean* = 1. Reciever Operating Characteristic (ROC) curves for the classification of nonoptimal codons defined as the bottom 25% of the tRNA adaptation index (tAI) (*59*) were computed with the R package pROC. Inhibitory codon pairs were defined as those in (*19*). For the comparison of consensus and mean RDs, their distributions were normalized to *mean* = 1 and *SD* = 1.

### Identification of differentially translated (DT) sequences

We focused on subsequences of length 3 codons as longer sequences rapidly drop in redundancy across the transcriptome thus limiting occurrences for further analysis. Differentially translated (DT) sequences were defined as sequences of 3 codons that had both at least 10 occurrences above gene-specific 90% and at least 10 occurrences below gene-specific 10% thresholds in their consensus profiles across all mRNAs. Thresholds were chosen as good trade-off between sufficient selectivity and data set size; subsequent results were found qualitatively very robust to these choices. For comparison, we extracted all sequences of length 3 codons with at least 20 occurrences throughout the yeast mRNAs. Sequences that were characterized by consistently high RD in the consensus (nDThi) were defined as those with at least 16 occurrences above the 90% threshold and at most 4 occurrences below the 10% threshold. Sequences that were characterized by consistently low RD in the consensus (nDTlo) were defined as those with at most 4 occurrences above the 90% threshold and at least 16 occurrences below the 10% threshold. Codon frequencies were compared to the background frequency of the mRNAs with sufficient coverage that passed our quality control (see above).

### Analysis of DT sequences

To test whether known sequence features that contribute to translation attenuation contribute to the observation of DT sequences, we analyzed a link to nonoptimal codon clusters, RNA folding strength, and clusters of positively charged amino acids. Nonoptimal codon clusters were defined as at least 2 nonoptimal codons in a stretch of length 5 translated within 2 codons, just before the current subsequence. Clusters of positively charged amino acids were defined as polypeptide stretches of length 8 amino acids containing at least 3 positive charges (Arg, Lys, His) with no negatively charged amino acids in between, and placed up to 35 sequence positions before, i.e. concurrently present in the ribosome exit tunnel. High RNA folding strength that can hamper ribosome translocation was tested by evaluating positive PARS scores (*107*) 4 codons downstream (*18*). Fisher’s exact test was used to test for statistical associations between the presence of one of these features and the category of fast and slowly translated subsequences. Domain boundaries of all yeast proteins were retrieved from (*108*), and binding sites of the yeast Hsp70 SSB were downloaded from (*32*). Average profiles of the sequences around SSB binding sites or domain boundaries were computed as per-position averages of the consensus RD profiles, and compared to averages of shuffled profiles of the defined regions. Gene ontology (GO) analyses were performed with the g:Profiler webserver (*109, 110*) through their Python API. All GO terms with a p-value of *p* < 0.001 for at least one input gene list are reported.

## ACKNOWLEDGMENTS

The authors are grateful for funding through a Discovery grant from the Natural Sciences and Engineering Research Council of Canada. S.P. holds the Canada Research Chair in Computational Systems Biology.

